# Ablation of B cell-derived IL-10 increases tuberculosis resistance

**DOI:** 10.1101/2024.07.17.603865

**Authors:** David Hertz, Sebastian Marwitz, Lars Eggers, Linda von Borstel, Gishnu Harikumar Parvathy, Jochen Behrends, Danny D. Jonigk, Rudolf A. Manz, Torsten Goldmann, Bianca E. Schneider

**Affiliations:** Host determinants in lung infections, Research Center Borstel, Leibniz Lung Center, Borstel, Germany; Core Facility Histology, Research Center Borstel, Leibniz Lung Center, Borstel, Germany; Airway Research Center North (ARCN), Member of the German Center for Lung Research (DZL), Großhansdorf, Germany; Core Facility Fluorescence Cytometry, Research Center Borstel, Leibniz Lung Center, Borstel, Germany; Institute of Pathology, University Hospital RWTH Aachen, Aachen, Germany; Biomedical Research In Endstage And Obstructive Lung Disease (BREATH), Member of the German Center for Lung Research (DZL), Hannover, Germany; Institute for Systemic Inflammation Research, University of Lübeck, Lübeck, Germany

## Abstract

Due to the historical dogma, that host defense against intracellular pathogens is mediated by cell-mediated immunity, B cells have been considered unimportant in providing protection against *Mycobacterium tuberculosis* (*Mtb*) and remained understudied for decades. However, emerging evidence suggests a more complex and multifaceted role for B cells in tuberculosis (TB) immunity. They accumulate at the side of infection in both animal models and human TB patients, suggesting a potential link to protective immunity. Still, the diverse roles of B cells in TB immunity continue to be unraveled. Apart from antibodies, B cells produce a wide range of cytokines, which can influence the local immune response. Here we addressed the relevance of interleukin 10 (IL-10) secreting B cells in long-term control of the *Mtb* Beijing strain HN878. Our research highlights the previously unknown role of B cell-derived IL-10 as a negative regulator of protective immunity in TB. For the first time, we demonstrate that mice lacking B cell-derived IL-10 show increased resistance to aerosol *Mtb* infection, as evidenced by a delayed onset of clinical symptoms and prolonged survival. Notably, this effect was significantly more pronounced in males compared to females, and was accompanied by male-specific immune alterations, indicating a previously unknown sex-specific regulatory role of B cell-derived IL-10 during *Mtb* infection.

## Introduction

Tuberculosis (TB) is the most prevalent bacterial infectious disease in humans with approximately 1.5 million deaths each year [1]. Gender- and sex-related factors lead to significant differences in prevalence between men and women [2–5], which is reflected by a male-to-female ratio for worldwide case notifications of 1.8 [1]. In a mouse model of TB - using either the reference *Mtb* strain H37Rv or the more virulent clinical isolate HN878 of the Beijing lineage - we previously found that males experienced a more rapid course of disease and succumbed to infection earlier than females, recapitulating the observed sex bias in human TB [6, 7]. Interestingly, higher male susceptibility to *Mtb* infection was associated with smaller B cell follicles in the lungs in comparison to *Mtb* infected females. B cell follicles are linked with immune protection in TB in mice, non-human primates, and humans [8–12]. Still, the underlying mechanisms and the functional role of B cells in TB remain poorly defined. A study by Swanson et al. recently showed that antigen-specific B cells direct T follicular-like helper cells into lymphoid follicles to mediate *Mtb* control [13]. However, the role of important B cell effector mechanisms such as antibody and cytokine production have yet to be elucidated.

Earlier studies have shown that the absence of B cells leads to heightened pulmonary inflammation and increased bacterial loads [14, 15], indicating that B cells play a protective, anti-inflammatory role in TB. B cells can suppress inflammation and modulate cellular immune responses by producing the anti-inflammatory cytokine interleukin-10 (IL-10). These so-called B10 cells are a sub-class of regulatory B-cells (Breg cells) and play a vital role in maintaining immune equilibrium by mitigating inflammation and averting autoimmune reactions [16]. Although IL-10 has beneficial effects in limiting tissue damage, its overproduction or dysregulated expression can favor *Mtb* persistence and chronic infection by limiting host-protective immune responses [17]. As such, T cell-derived IL-10 was shown to contribute to IL-10-induced susceptibility to TB [18]. However, the role of B cell-derived IL-10 is still not fully understood and requires further investigation. B10 cells were detected in *Mtb* infected mouse lungs, especially during the chronic phase [18] and in blood of patients with active TB [19]. *In vitro* stimulation assays confirmed that B cells produce IL-10 in response to mycobacterial antigen [20, 21]. Mannose-capped lipoarabinomannan (ManLAM), a major surface lipoglycan component from *Mtb*, was shown to be a very potent inducer of B10 cells *in vitro* and *in vivo* [19]. Notably, these B10 cells inhibited Th1 polarization, and promoted Th2 polarization, suggesting that ManLAM-induced B10 cells negatively regulate anti-*Mtb* immunity. In order to gain a clearer understanding of the precise involvement of B10 cells in TB, we opted to delve deeper into their functionality *in vivo*. To do so, we infected male and female mice harboring a targeted knockout of IL-10 in B cells (IL-10^flox^/CD19^cre^) with *Mtb* HN878 and monitored disease progression, control of bacterial replication and immunological changes over the course of infection. Absence of B cell-derived IL-10 significantly improved survival of HN878 infected mice compared to their B cell IL-10 competent littermates, but of note, the survival advantage was higher in males compared to females. Improved male survival was associated with reduced bacterial loads and profound alterations in the lung immune environment. Collectively, our findings provide compelling evidence that B cell-derived IL-10 plays a far more critical role in regulating the male immune response to *Mtb* infection compared to females. These findings highlight the critical need for further investigations into the mechanisms underlying sex-based differences in TB immunity.

## Material and Methods

### Mice

C57BL/6j mice were bred under specific-pathogen-free (SPF) conditions at the Research Center Borstel. IL-10 reporter mice (Vert-X) [22] were bred under SPF conditions at the animal facility of the University of Lübeck. IL-10 flox/flox mice (B6.129P2(C)-Il10tm1Roer/MbogJ), with loxP sites that flank exon 1 of IL-10, were bred under SPF conditions with CD19-Cre mice (B6.129P2(C)-Cd19tm1(cre)Cgn/J) at the animal facility of the University of Lübeck, resulting in the generation of mice with B cell-specific inactivation of IL-10 (B cell IL-10 KO), and IL-10 competent littermate controls [23]. For infection experiments, male and female mice aged between 11-20 weeks were maintained under barrier conditions in the BSL 3 facility at the Research Center Borstel in individually ventilated cages. Animal experiments were in accordance with the German Animal Protection Law and approved by the Ethics Committee for Animal Experiments of the Ministry of Agriculture, Rural Areas, European Affairs and Consumer Protection of the State of Schleswig-Holstein.

### *Mtb* infection and determination of bacterial load

*Mtb* HN878 was grown in Middlebrook 7H9 broth (BD Biosciences, Franklin Lakes, USA) supplemented with 10% v/v OADC (Oleic acid, Albumin, Dextrose, Catalase) enrichment medium (BD Biosciences, Franklin Lakes, USA) to logarithmic growth phase (OD_600_ 0.2 - 0.4) and aliquots were frozen at -80°C. Viable cell number in thawed aliquots were determined by plating serial dilutions onto Middlebrook 7H11 agar plates supplemented with 10% v/v OADC followed by incubation at 37°C for 3-4 weeks. For infection of experimental animals, *Mtb* stocks were diluted in sterile distilled water at a concentration providing an uptake of approximately 50 viable bacilli per lung. Infection was performed via the respiratory route by using an aerosol chamber (Glas-Col, Terre-Haute, IN, USA) as described previously [6]. The inoculum size was quantified 24 h after infection in the lungs and bacterial loads in lung and spleen were evaluated at different time points after aerosol infection by determining colony forming units (CFU). Organs were removed aseptically, weighed, and homogenized in 0.05% v/v Tween 20 in PBS containing a proteinase inhibitor cocktail (Roche, Basel, Switzerland) prepared according to the manufacturer’s instructions for subsequent quantification of cytokines (see below). Tenfold serial dilutions of organ homogenates in sterile water/1% v/v Tween 80/1% w/v albumin were plated onto Middlebrook 7H11 agar plates supplemented with 10% v/v OADC and incubated at 37°C for 3–4 weeks.

### Clinical Score

Clinical score was used to indicate severity of disease progression [7]. Animals were scored in terms of general behavior, activity, feeding habits, and weight gain or loss. Each of the criteria is assigned score points from 1 to 5 with 1 being the best and 5 the worst. The mean of the score points represents the overall score for an animal. Animals with severe symptoms (reaching a clinical score of ≥3.5) were euthanized to avoid unnecessary suffering, and the time point was recorded as the end point of survival for that individual mouse.

### Flow Cytometry

Mice were sacrificed and perfused intracardially with sterile PBS before organ harvest. Lungs were digested in 100 µg/ml DNase I (Roche) and 50 µg/ml Liberase TL (Roche) in RPMI for 60 minutes and passed through a 100 µm pore size cell strainer to obtain a single cell suspension. Single-cell suspensions were blocked with anti-CD16/CD32 (clone 93, Biolegend, Fell, Germany) and sera (rat, rabbit, mouse (PAN Biotech, Aidenbach, Germany) and hamster (abcam, Cambridge, UK)) for 30 min at 4°C. Subsequently, cells were washed and incubated for 30 min at 4°C with the following antibodies: BV510 anti-CD45 (clone 30-F11); BV421 or PerCP-Cy5.5 anti-CD138 (clone 281-2); APC anti-GL7 (clone GL7) from BioLegend, Fell, Germany and PE-Cy7 anti-B220 (clone RA3-6B2) from BD Biosciences, Franklin Lakes, USA. After washing, cells were resuspended in FACS buffer (3% fetal calf serum heat-inactivated, 0.1 % NaN_3_, 2 mM EDTA) and analyzed on a MACSQuant Analyzer 10 (Miltenyi Biotec B.V. & Co. KG, Bergisch Gladbach, Germany). Imaging of IL-10 producing B cells were performed on a BD FACSDiscover™ S8 Cell Sorter with BD CellView^™^ Image Technology and BD SpectralFX^™^ Technology (BD Biosciences, Franklin Lakes, USA). Data were analyzed with the FCSExpress software (DeNovo^TM^ Software, Pasadena, CA, USA) and the Cytolution software (Cytolytics GmbH, Tübingen, Germany).

### Multiplex cytokine assay

The concentrations of various cytokines and chemokines in lung homogenates were determined by LEGENDplex^TM^ (Mouse Inflammation panel and Mouse Proinflammatory Chemokine Panel, BioLegend, Fell, Germany) according to the manufacturer’s protocol.

### Histology

Superior lung lobes from infected mice were fixed with 4% w/v paraformaldehyde for 24 h, embedded in paraffin, and sectioned (4 μm). Sections were stained with hematoxylin and eosin (H&E). Slides were imaged with a PhenoImager™ HT instrument (Akoya Biosciences, Marlborough, USA). The quantitative analysis of inflammatory area was conducted by determination of whole lung area and inflammatory area for H&E-stained sections using the software QuPath (https://qupath.github.io/) [24].

### Multiplex-immunofluorescence (mIF) staining

Formalin-fixed, paraffin-embedded (FFPE) samples of mouse lung tissues were deparaffinized by immersion into xylene (2x 10 min) and rehydrated by graded series of ethanol (2x 2 min 100%, 2 min 96%, 2 min 90%, 2 min 80%, 2 min 70%, 2 min distilled water). Rehydrated samples were transferred to 1x TBST (50 mM Tris, pH 7.6, 0.05% Tween-20) and subsequently forwarded to mIF staining. The Opal 3-Plex Anti-Rb Manual Detection Kit (Akoya Biosciences, Marlborough, USA, NEL840001KT) was used with additional fluorophores as indicated in Supplementary Table S1. Overall, mIF staining consisted of several, subsequent cycles of IF staining targeting one antigen per time: Before antigen detection, endogenous peroxidases were blocked once by incubation for 10 min at RT in 3% H_2_O_2_ followed by washing with 1x TBST. One IF cycle consisted of 10 min protein blocking using antibody diluent (Akoya Biosciences, Marlborough, USA) followed by 3x 2 min washing with 1x TBST. Primary antibodies were diluted using antibody diluent (Akoya Biosciences, Marlborough, USA) and incubated for 45 min at RT followed by 3x 3 min washing with 1x TBST. Primary antibodies were detected by incubation for 10 min with HRP-Polymer (Akoya Biosciences, Marlborough, USA) and subsequent TBST washing for 3x 3 min. Signals were visualized by OPAL-TSA reaction for 10 min at RT and stopped by washing 3x for 3 min with 1x TBST. An antigen-retrieval (AR) step by incubation in citric acid buffer pH6 (Akoya Biosciences, Marlborough, USA) for 1 min at 1000 W and 10 min at 100 W was used to remove primary antibodies and HRP polymer before entering the next staining cycle. Finally, nuclei were stained using spectral DAPI (Akoya Biosciences, Marlborough, USA) and slides were mounted with cover slips using ProLong Gold (Invitrogen, Carlsbad, USA). Please see Supplementary Table S1 for a detailed pipetting protocol for the mIF panel. The panel was developed using positive control tissues and antigen stability was tested for each antibody by increasing rounds of AR using citric acid buffer pH6 to determine optimal positioning within each mIF panel. Single-plex stains for each target of the multiplex panel from the respective cycle were conducted and their results qualitatively compared to multiplex results for concordance of overall staining pattern, intensity and frequency of detected cells.

### mIF Imaging & digital image analysis

Whole-slide imaging (WSI) was performed using a PhenoImager HT multi-spectral slide scanner (Akoya Biosciences, Marlborough, USA) at 20x magnification (0.5 µm/pixel) with exposure times held constant for all analyzed samples (DAPI: 1.39ms, Opal 480: 10 ms, Opal 520: 20 ms, Opal 570: 55 ms, Opal 780: 15 ms). Phenochart software 1.1.0 (Akoya Biosciences, Marlborough, USA) was used to select regions of interest for generation of image analyses algorithm as well as batch analyses across all tissues. For this, an analysis algorithm was built using InForm software 2.6.0 (Akoya Biosciences, Marlborough, USA) with representative regions across different samples. The image analysis algorithm consisted of several steps including spectral unmixing based on proprietary spectral libraries of designated fluorochromes as supplied by Akoya Biosciences, Marlborough, USA, pixel-based tissue segmentation, cell segmentation and cell classification. Representative regions from several different WSIs were imported into InForm software, spectrally unmixed and forwarded to user-guided, machine-learning tissue segmentation based on manual annotation for areas consisting either of empty glass/no tissue, B cell enriched tissue and tissue without B cell enrichment. Tissue segmentation accuracy was tested on randomly selected WSIs and re-trained on areas where initial tissue segmentation failed to correctly identify designated areas. Once the tissue segmentation reached satisfactory level, individual cells were segmented based on their DAPI signal and accessory information from antibody staining. A cell classification algorithm was trained to identify each antigen in a binary approach (antigen positive vs negative). For analysis of WSIs, the complete tissue was deconstructed into boxes and subjected to batch analyses. All images were individually reviewed and areas containing imaging artifacts (dust, fibres, tissues folds, out-of-focus) as well as pleura with excess autofluorescence removed. Resulting cell segmentation files of 828 images from 18 WSIs were merged and consolidated into a single data file using R package phenoptr reports 0.3.3 [25]. Phenoptr reports was further used to determine the cell density (cells/mm^2^) for selected combinations of detected antigens across the complete tissue of each animal. The slides were analyzed in a blinded manner.

### NanoString analysis

Superior lung lobes from infected mice were fixed with 4% w/v paraformaldehyde for 24 h and embedded in paraffin. RNA was extracted with RNeasy FFPE Kit (Qiagen, Germantown, USA) from tissue sections, dissolved in nuclease-free water and RNA integrity analyzed using a 2100 Bioanalyzer (RNA 6000 Nano Kit, Agilent Technologies, Waldbronn, Germany). Samples with DV200 values >30% were included for gene expression analysis. The NanoString nCounter (NanoString Technologies, Inc., Seattle WA, USA) platform was used to assess the effects of B cell derived IL-10 on changes in immune gene expression. Briefly, RNA (n = 3/group) was analyzed using the nCounter Host response panel (catalog # 115000486), which includes 773 immune-related genes and 12 internal reference genes for data normalization. Assays were performed using a nCounter Prep Station 5s (NanoString Technologies, Inc., Seattle WA, USA) according to the manufacturer’s instructions, and the proceeded cartridge was sent for digital analysis using the nCounter® Sprint Profiler system (performed at the Institute of Pathology, Hannover Medical School, Hannover, Germany). Briefly, RNA is directly tagged with a capture probe and a reporter probe specific to the genes of interest. After hybridization, samples were washed and the purified target-probe complexes were aligned and immobilized onto the nCounter cartridge. The transcripts were counted via detection of the fluorescent barcodes within the reporter probe. Raw gene expression data were analyzed using NanoString’s software nSolver v4.0 with the Advanced Analysis Module v2.0.134 and ROSALIND® (https://nanostring.rosalind.bio).

### Statistical Analysis

All data were analyzed using GraphPad Prism 10 (GraphPad Software, La Jolla, USA) or Excel 2016 (Microsoft Corporation, Redmond, USA). Statistical tests are indicated in the individual Figure legends. Correlation was determined by calculating Pearson’s coefficient using a two-tailed analysis. A correlation was taken into account as of r≥ 0.60 (defined as strong correlation). Principal Component Analysis (PCA) was performed in BioVinci version 3.0.9 (BioTuring INc., San Diego, USA) – a machine learning aided analysis platform for biological datasets.

## Results

### B cells contribute to IL-10 production after Mtb infection

It has been previously shown that B cells are a source of IL-10 during *Mtb* HN878 infection in female mice [18]. Since differences in the frequencies of IL-10-producing B cells have been described between the sexes in mouse models of arthritis [26], we assessed B cell-specific IL-10 expression after infection with *Mtb* in a sex-specific manner. To do so, male and female IL-10 transcriptional reporter (Vert-X) mice which express enhanced green fluorescent protein (eGFP) under the control of the IL-10 promoter [22] were infected via the aerosol route with *Mtb* HN878 (Fig. 1; gating strategy see Supplemental (Suppl.) Fig. S1). At the indicated time points, single cell suspensions of lung tissues were prepared and B cells were analyzed for the expression of GFP (Fig. 1A). IL-10 expressing B cells, defined as GFP^+^CD138^-^B220^+^ B cells and GFP^+^CD138^+^B220^var^ plasma B cells, respectively, were detected in the lungs of both sexes at all-time points investigated. Even though the proportion of these cells within the overall population of IL-10 producers remained constant (Fig. 1B), the total number of IL-10-producing B cells increased over time (Fig. 1C). While UMAP analysis revealed a consistent increase in the proportion of GL7^+^ germinal center B cells, CD138^+^ plasma cells (PCs) remained a relatively small population throughout the infection (Fig. 1D and 1E). However, PCs were by far the most significant producers of IL-10 at all-time points analyzed (Fig. 1F-1J). This aligns with established literature that identifies PCs as the major source of B cell-derived IL-10 *in vivo* across various disease models, including autoimmune, malignant and infectious conditions [27]. Statistical analysis revealed that the proportion of IL-10^+^ PCs was significantly higher in males compared to females early after *Mtb* infection (d13; Fig. 1H), without significant differences in the expression of IL-10 on a per cell basis (Fig. 1I).

**Figure 1.**
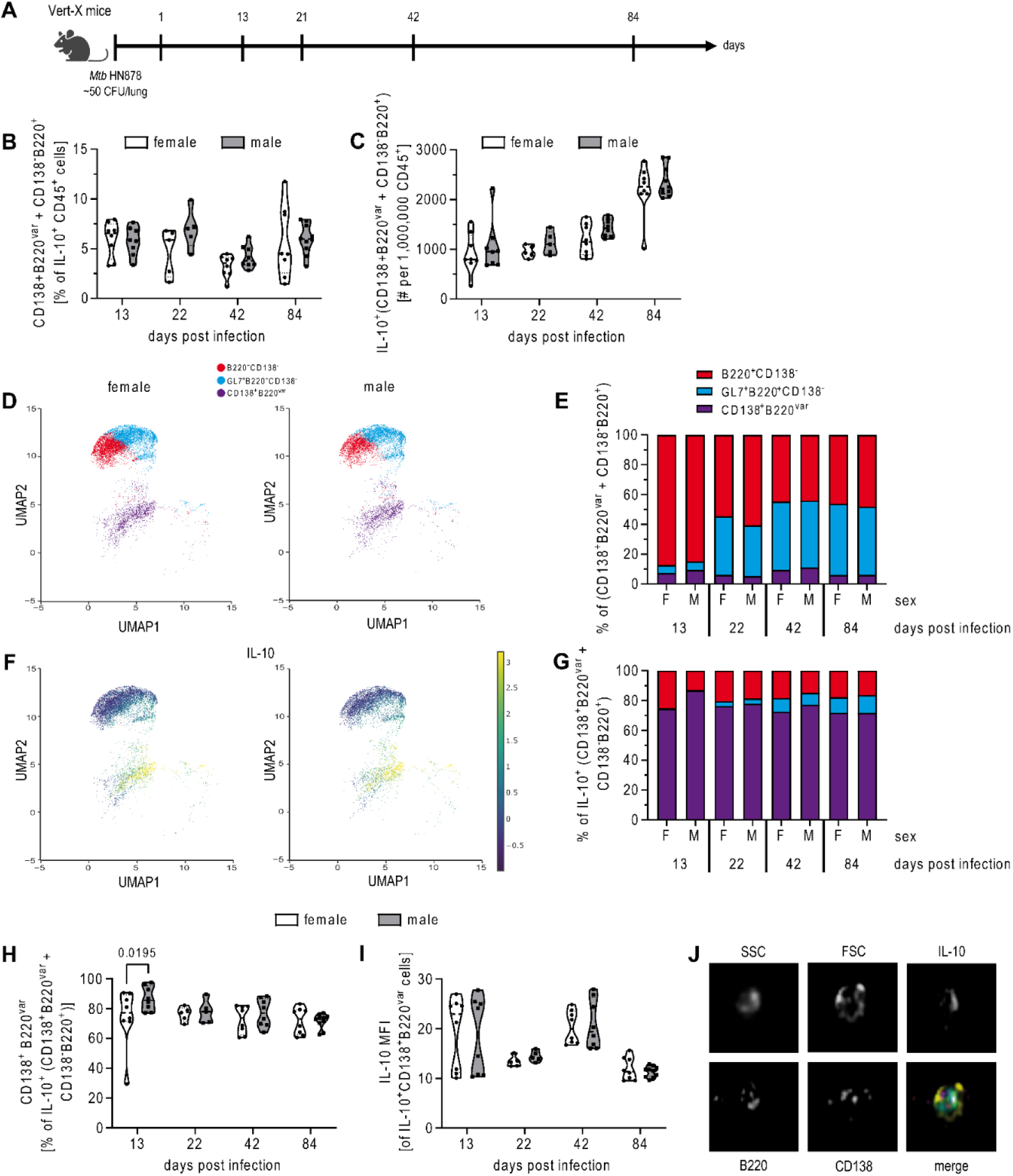
B cells produce IL-10 after Mtb infection. A) Experimental setup. Female and male IL-10 reporter mice (Vert-X) were infected via aerosol with *Mtb* HN878 and IL-10 producing B cells in the lung were determined at the indicated time points by flow cytometry. B and C) Total B cells (defined as CD138^+^B220^var^ + CD138^-^B220^+^) as percentage of IL-10^+^CD45^+^ cells and as total number of one million CD45^+^ cells. D and E) Plasma cells (CD138^+^B220^var^), germinal center B cells (GL7^+^CD138^-^B220^+^) and other B cell subtypes (CD138^-^B220^+^) as percentage of total B cells. F and G) IL-10 producing B cell subtypes as percentage of total IL-10^+^ B cells. H) Plasma cells as percentage of total IL-10^+^ B cells and I) median fluorescence intensity (MFI) of IL-10 in IL-10 producing plasma cells. J) IL-10^+^ plasma cell of a female Vert-X mouse from day 42 p.i.. B, C, H and I: Each data point represents one mouse. Data were combined from 2 independent experiments; n = 5-10. Statistical analysis was performed by 2way ANOVA followed by Tukey’s multiple comparisons test. D and F: Data were generated from 4 female and 5 male mice on day 84 p.i. from 1 representative out of 2 experiments. E and G: Data are represented as the mean percentage of B cell subtypes to total B cells from 2 independent experiments; n=5-10.

Overall, B cells contribute to IL-10 production in the *Mtb* infected lung in both sexes, with CD138^+^ PCs representing the primary source of B cell-derived IL-10.

### IL-10 deficiency in B cells renders mice more resistant to Mtb HN878 infection

To determine the actual role of B cell-derived IL-10 during *Mtb* infection we evaluated disease progression in mice that were deficient in IL-10 expression specifically in B cells (*Cd19-cre^+/-^Il10^fl/fl^;* B cell IL-10 KO) (Fig. 2A). B cell IL-10 KO mice were more resistant to the *Mtb* infection than their littermates as reflected by reduced body weight loss and a prolonged survival (Suppl. Fig. S2A and Fig. 2B). Intriguingly, segregation of survival data by sex revealed that the survival benefit in the absence of B cell-derived IL-10 was more pronounced in males compared to females (Fig. 2C and 2D). Together, these data clearly argue against a protective role for B cell-derived IL-10 but instead suggest it increases susceptibility to TB, particularly in males.

**Figure 2.**
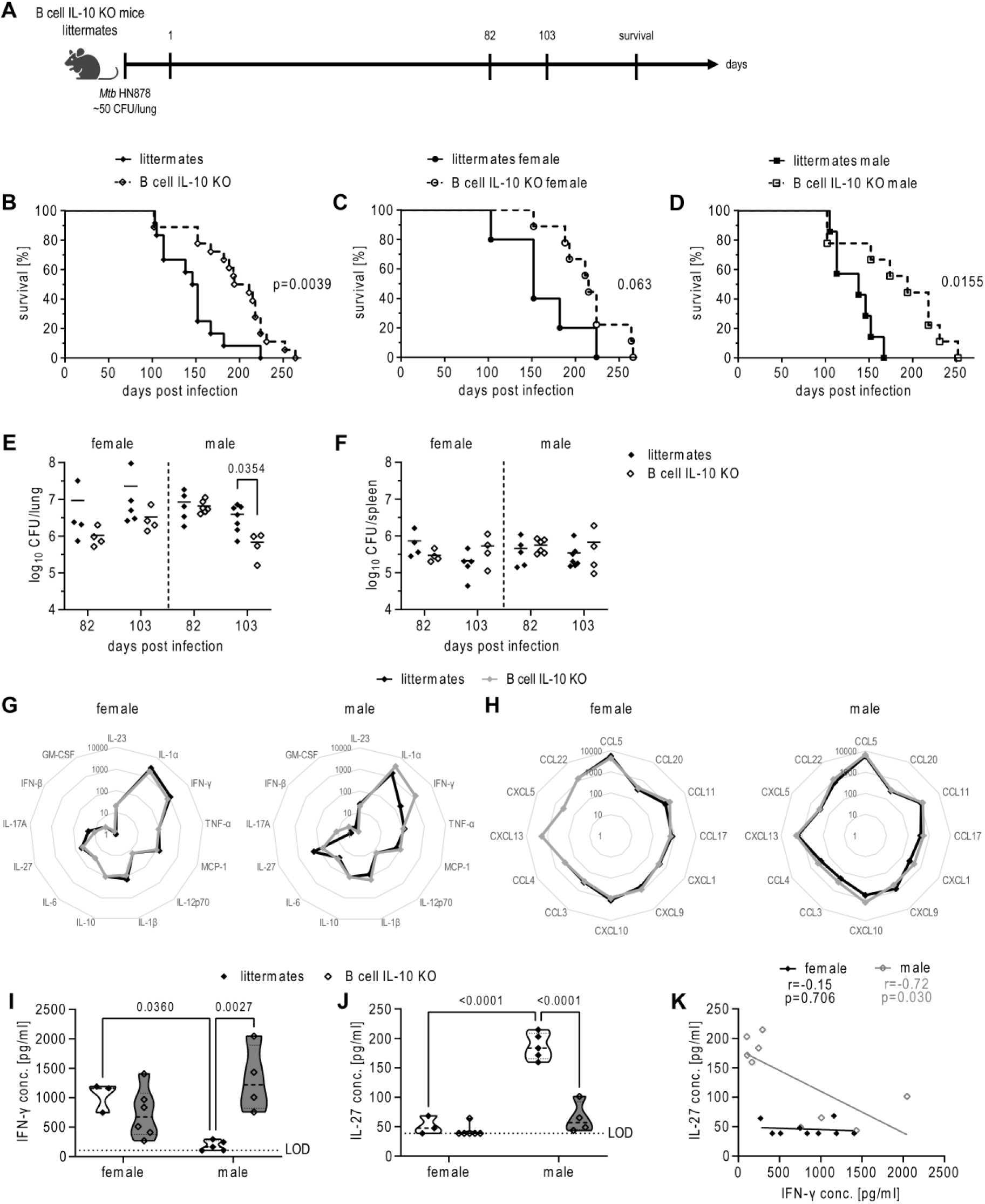
Ablation of B cell derived IL-10 increases resistance of mice to Mtb infection. A) Experimental setup. Female and male B cell IL-10 KO mice and their B cell IL-10 competent littermates were infected via aerosol with *Mtb* HN878 and analyzed at the indicated time points. B) Survival of female (n=9) and male (n=9) B cell IL-10 KO mice and their female (n=5) and male (n=7) B cell IL-10 competent littermates. Survival stratified by sex is shown in (C) for females and (D) for males. CFU of the lung (E) and spleen (F) for female and male littermates and B cell IL-10 KO mice at day 82 p.i. and day 103 p.i. Spider chart of cytokines (G) and chemokines (H) measured in lung homogenates of female and male littermates and B cell IL-10 KO mice at day 82 p.i. Concentrations of cytokines/chemokines are shown as pg/ml. IFN-γ (I) and IL-27 (J) in lung homogenates from female and male littermates and B cell IL-10 KO mice at day 82 p.i. K) Correlation of IFN-γ and IL-27 levels as pooled data for littermates and B cell IL-10 KO mice for both sexes. B-D) Each data point represents one mouse from one experiment. Statistical analysis was performed by log rank test. E and F) Each data point represents one mouse from one representative experiment out of two (n = 4-7). Statistical analysis was performed by Student’s t-test. G-K) Each data point represents one mouse from one representative experiment out of two (n = 3-6). Statistical analysis was performed by 2way ANOVA followed by Tukey’s multiple comparisons test. Correlation was calculated using Pearson correlation.

### Ablation of B cell IL-10 improves control of Mtb and alters the cytokine profile in male lungs

We next investigated whether the improved survival observed in mice lacking B cell-derived IL-10 resulted from a better ability to control bacterial loads after HN878 infection. Day 82 analysis of lung tissue revealed no significant difference in bacterial burden between groups. However, by day 103, male B cell IL-10 KO mice exhibited a significantly lower lung bacterial load compared to male littermates (Fig. 2E), who displayed initial signs of disease around this time (Suppl. Fig. S2C). B cell IL-10 deficiency did not affect bacterial burden in the spleen (Fig. 2F), suggesting distinct roles for B cell-derived IL-10 in the lung and spleen during *Mtb* infection.

To examine the immunological environment in the lungs that accompanied improved bacterial control in the absence of B cell IL-10, we quantified the levels of inflammatory cytokines and chemokines known to play a role in *Mtb* infection. While absence of B cell IL-10 did not lead to overall changes in the levels of cyto- or chemokines in female lungs (Fig. 2G and 2H), it had profound effects on certain cytokines in male lungs (Fig. 2G). Specifically, we observed significant changes in IFN-γ and IL-27 levels, with differences being evident not only between littermate and B cell IL-10 KO mice, but also between males and females (Fig. 2I and 2J). Interestingly, male littermates displayed significantly lower levels of IFN-γ compared to females (Fig. 2I), which increased significantly in the absence of B cell-derived IL-10. In contrast to IFN-γ, IL-27 levels were significantly higher in male littermates compared to females (Fig. 2J). Notably, the absence of B cell IL-10 led to a significant decrease in IL-27 levels specifically in male mice. This reduction brought their IL-27 levels down to a level comparable to that observed in female mice. This difference in how B cell IL-10 regulates IFN-γ and IL-27 between the sexes is further supported by correlation analysis (Fig. 2K). A strong negative correlation between IL-27 and IFN-γ levels was observed in male mice, while this negative correlation was absent in female mice, indicating a possible different regulatory pathway for these cytokines in female mice compared to male mice.

### Histopathological changes in lung tissue from Mtb infected B cell IL-10 KO mice

IL-27, like IL-10, is an important anti-inflammatory cytokine that has a dual role in TB. On the one hand, IL-27 limits protective immunity to *Mtb* but on the other hand it prevents from excessive inflammation and immunopathology [28]. Thus, we next evaluated the granulomatous response in absence of B cell-derived IL-10, which led to a significant reduction in IL-27 levels in males (see Fig. 2J). To this end, H&E-stained lung sections from B cell IL-10 KO mice and littermates were examined 82 days after HN878 infection (Fig. 3A). No significant difference in cellular infiltration was observed in the lungs of female B cell IL-10 KO mice compared to their female littermates. In contrast, male B cell IL-10 KO mice exhibited a greater degree of cellular infiltration compared to their male littermates (Fig. 3B). In fact, the absence of B cell-derived IL-10 in males led to levels of cellular infiltration in the lungs comparable to those observed in females. This suggests that B cell-derived IL-10 plays a specific role in down-regulating the inflammatory response within the lungs of male mice during *Mtb* infection. The results from the histological examination align remarkably well with the cytokine analysis, which is further supported by the observed correlations (Fig. 3C and 3D). In males, a strong positive correlation exists between the area of inflammation and IFN-γ levels (Fig. 3C), while conversely, a negative correlation was observed between inflammation and IL-27 levels (Fig. 3D). Notably, these correlations were absent in females, indicating a potentially distinct inflammatory pathway in females that is not influenced by B cell-derived IL-10.

**Figure 3.**
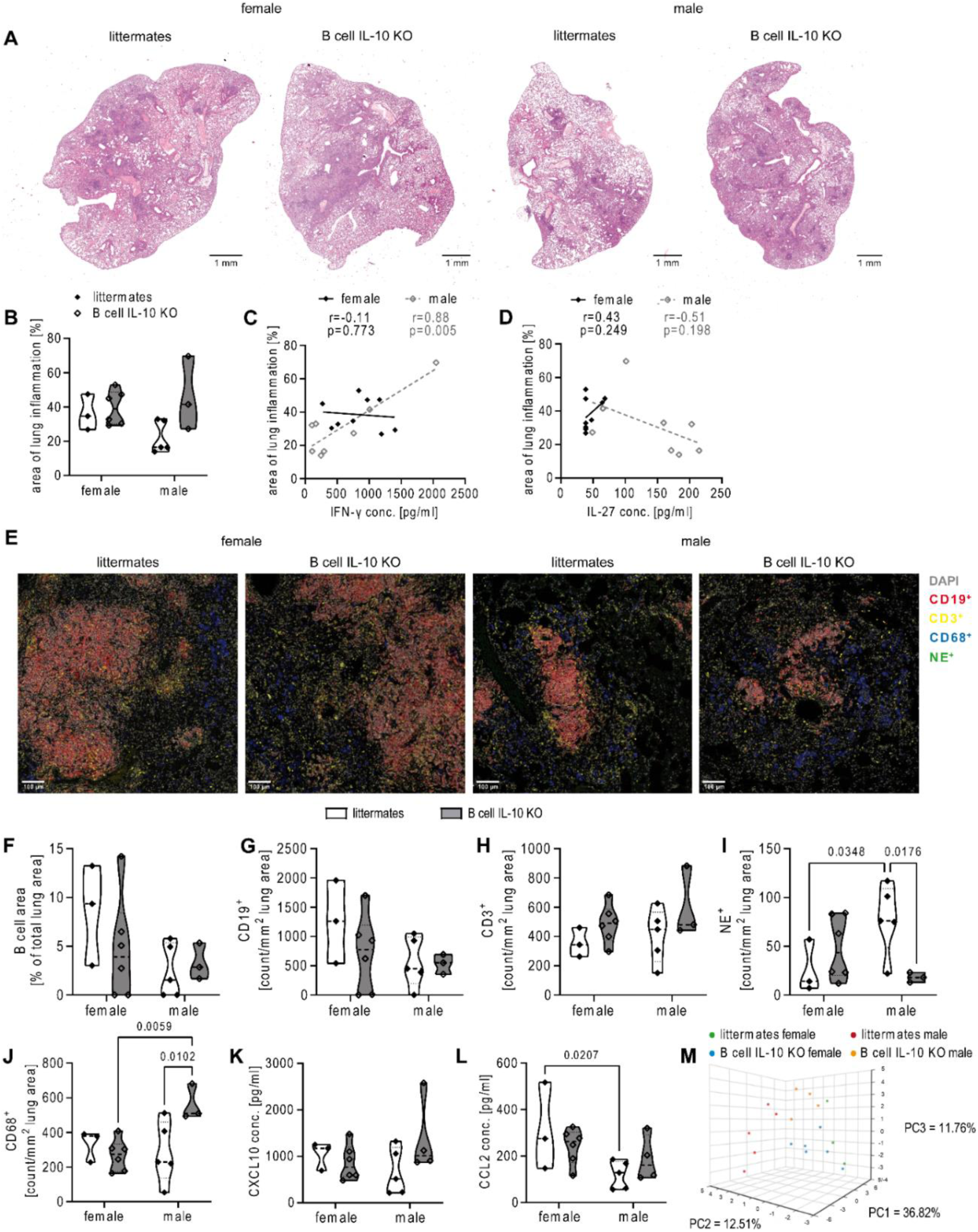
Ablation of B cell IL-10 increases cellular infiltration into the male lung. Female and male B cell IL-10 KO mice and B cell IL-10 competent littermates were infected via aerosol with *Mtb* HN878. Lungs were collected at day 82 and PFA-fixed, paraffin embedded tissue sections were stained with H&E. A) Representative micrographs from one female and male mouse lung per genotype are shown. Bar = 1 mm. B) Quantitative analysis of the area of lung inflammation shown in (A) segregated by sex and correlations with the levels of IFN-γ (C) and IL-27 (D). E) Representative micrographs from lungs of one female and male mouse per group stained with antibodies to detect B cells (CD19^+^), T cells (CD3^+^), neutrophils (NE^+^) and macrophages (CD68^+^); Bar = 100 µm. F-J) Quantitative analysis of B cell area and respective immune cells as shown in (E). Levels of CXCL10 (K) and CCL2 (L) in lung homogenates from female and male littermates and B cell IL-10 KO mice at day 82 p.i. M) PCA of integrated datasets (Suppl. Table S2). Datasets were standardized using the default algorithm in BioVinci and integrated into a PCA plot. B-D and F-M) Each data point represents one mouse from one representative experiment out of two; n = 3-6. Statistical analysis was performed by 2way ANOVA followed by Tukey’s multiple comparisons test (B; F-L). Correlation was calculated using Pearson correlation (C and D).

The sex-specific differences in cellular infiltration and the potential role of B cell-derived IL-10 in males prompted a more detailed examination of immune cell populations within the infected lung using mIF to identify CD19^+^ B cells, CD3^+^ T cells, CD68^+^ macrophages, and neutrophil elastase positive (NE^+^) neutrophils (Fig. 3E). Consistent with our previous findings [7], the total area and density of CD19^+^ B cells within the lesions were lower in male compared to female littermates, although not statistically significant (Fig. 3F and 3G). Interestingly, B cell IL-10 deficiency did not significantly affect B cell area or density in either sex. Likewise, our analysis of CD3^+^ T cells revealed no major difference between littermates and B cell IL-10 KO mice, or – unlike CD19^+^ B cells - between the sexes (Fig. 3H). In contrast, B cell IL-10 deficiency significantly impacted macrophages and neutrophils in males. Male littermates exhibited a significantly higher density of NE^+^ neutrophils within their lesions compared to females, and B cell IL-10 deficiency led to a dramatic reduction in neutrophils specifically in males, but not in females (Fig. 3I). Conversely, male B cell IL-10 KO mice had a marked increase in CD68^+^ macrophages compared to littermates, while macrophage density in females remained similar regardless of B cell IL-10 status (Fig. 3J). Consistent with the histological data, we found a trend towards more CXCL10 and CCL2 in male but not in female B cell IL-10 KO mice (Fig. 3K and 3L).

In order to understand the combined influence of the measured parameters described, we employed Principal Component Analysis (PCA) on our comprehensive dataset of day 82 post infection (Suppl. Table S2). The first three principal components (PCs) captured a significant portion (61%) of the total variance within the dataset (Fig. 3M). The PCA plot revealed a distinct separation between male littermates and all other groups. Interestingly, male B cell IL-10 KO clustered closer to the combined group of female littermates and female B cell IL-10 KO mice. This suggests a sex-based difference in the overall immune response to *Mtb* infection. Moreover, the closer proximity of male B cell IL-10 KO mice to females aligns with their improved survival compared to male littermates.

Overall, these data suggest that the absence of B cell-derived IL-10 has a more pronounced effect on the immune response in males, making them more susceptible to *Mtb* in its presence.

### Gene expression analysis reveals B cell IL-10 dependent regulation of immune response gene transcripts in a sex-specific manner

To better understand the molecular basis of the observed sex differences in susceptibility and lung inflammation following *Mtb* infection in mice that lack B cell-derived IL-10, we employed the nCounter platform to analyze immune-related gene expression profiles in the lungs that we had analyzed by mIF before. Sex-wise gene expression analysis revealed a striking disparity: Only few genes were differentially expressed in female B cell IL-10 KO mice compared to female controls (Fig. 4A). In stark contrast, male B cell IL-10 KO mice displayed significant changes, with 114 genes being differentially regulated, most of which were downregulated (Fig. 4B). Quantitative analysis of pathway scores revealed a striking sex difference in response to B cell IL-10 deficiency. Pathway scores in males showed greater variation compared to their female counterparts, where scores remained relatively unchanged by B cell IL-10 depletion (Fig. 4C and 4D). Notably, the JAK-STAT pathway, known to play a central role in mediating the effects of IL-10, exhibited the most significant downregulation in male B cell IL-10 KO mice (Fig. 4E). Additionally, pathways related to B cell receptor (BCR) and IL-2 signaling were also downregulated in males, while the host defense peptides pathway showed a contrasting increase (Fig. 4F-H).

**Figure 4.**
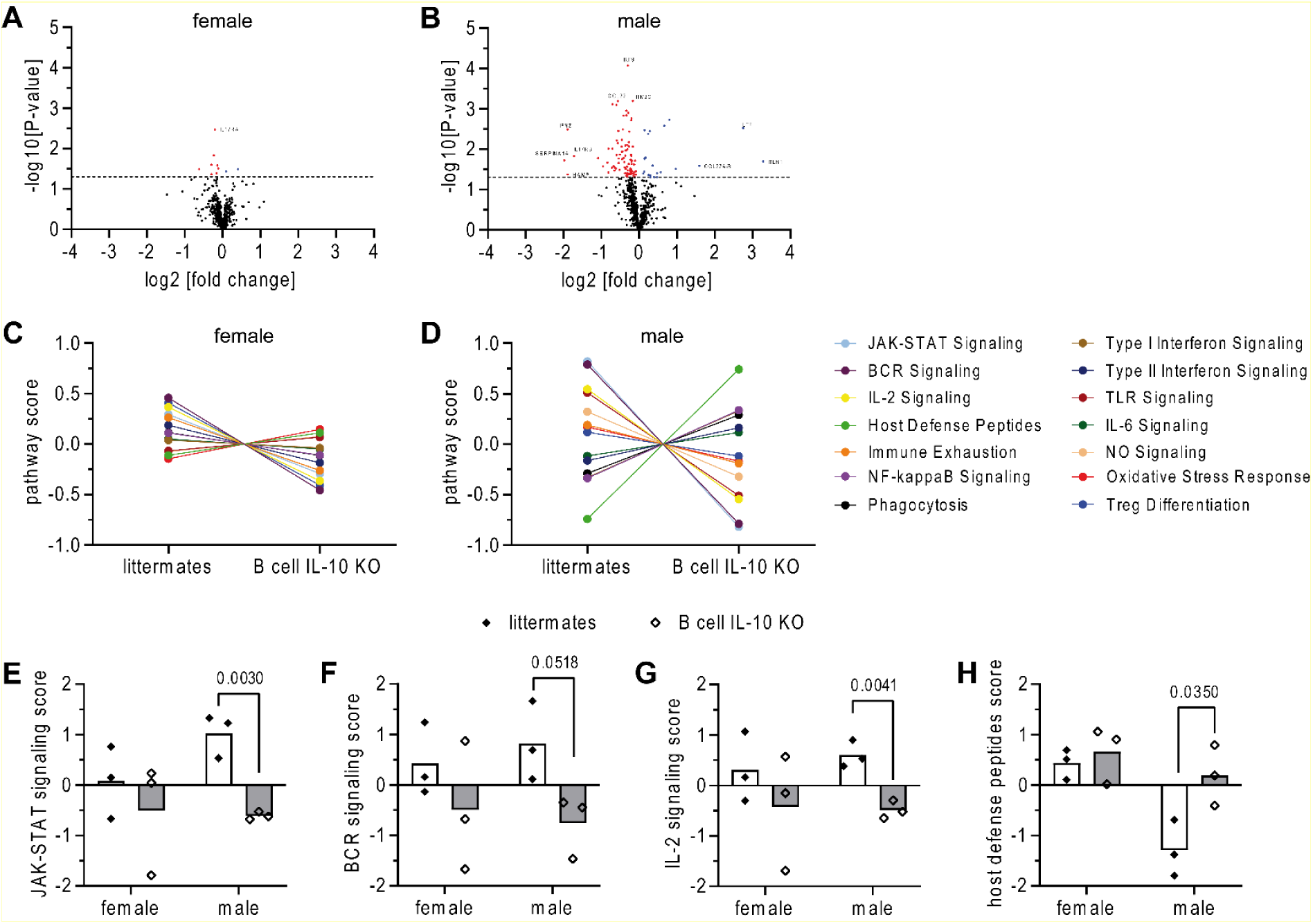
Ablation of B cell derived IL-10 leads to significant changes in immune response gene transcripts in a sex-specific manner. B cell IL-10 KO mice and their B cell IL-10 competent littermates were infected via aerosol with HN878. Lungs were collected at day 82 and PFA-fixed, paraffin embedded tissue sections were used to extract RNA. Gene expression was analyzed with the nCounter Host Response Panel by Nanostring technology. A and B) Volcano plots of log2 fold change versus - log10 p-value of the differential gene expression between female (A) and male (B) B cell IL-10 KO mice and their B cell IL-10 competent littermates. Horizontal dot line represents p-value <0.05. Significantly upregulated genes in B cell IL-10 KO mice are represented as blue dots, significantly downregulated genes are red; n = 3/group. C and D) Selected trend plots of pathway signatures for female and male littermates and B cell IL-10 KO mice. E-H) Most significantly different pathway signatures are depicted for female and male B cell IL-10 KO mice and their B cell IL-10 competent littermates. Each data point represents one mouse from one experiment; n = 3/group. Statistical analysis was performed by Student’s t-Test.

Collectively, these findings provide compelling evidence for our hypothesis that B cell-derived IL-10 plays a far more critical role in regulating the male immune response during *Mtb* infection compared to females.

## Discussion

While IL-10-producing B cells have been identified in *Mtb* infection across mice and humans [18–21], their presence and function have not been investigated from a sex-specific perspective. Moreover, earlier studies suggested that B cells participate in *Mtb* control independent of the production of IL-10 [13] while T cells have been recognized as the major source of IL-10 that promotes TB susceptibility [18]. These earlier studies demonstrated that B cell IL-10 deficient mice could still control *Mtb*. However, Moreira-Teixeira used only female mice and evaluated bacterial growth after 60 days of infection [18], while Swanson determined bacterial burden 50 and 100 days post infection but did not report on the sex of the mice [13]. In contrast to these studies, we investigated the impact of B cell IL-10 in both male and female mice and monitored their survival in addition to bacterial burden. In line with the previous results, we also did not observe a reduction in bacterial load in the absence of B cell IL-10 in females but in males, and these mice also exhibited significantly improved survival.

Our study reveals a previously unidentified role for B cell-derived IL-10 in regulating immunity during TB. Intriguingly, immunological analysis, including gene expression data using NanoString technology, demonstrated a significantly broader impact of B cell IL-10 on the male immune response compared to females. While female B cell IL-10 KO mice displayed minimal changes in cytokine/chemokine profiles or gene expression, ablation of B cell IL-10 had a profound effect on the immunological environment in the male lung. Notably, it reversed the sex-specific pattern of IFN-γ and IL-27 which likely contributes significantly to the higher susceptibility of males to *Mtb* infection. Male control mice exhibited significantly lower IFN-γ but higher IL-27 levels compared to females. IFN-γ is essential to control TB infection [29, 30], whereas IL-27 has been previously shown to exhibit a double-edged sword effect: suppressing protective immunity against TB and bacterial clearance, while simultaneously dampening excessive inflammation [28, 31, 32]. Studies using mice deficient in WSX-1, a component of the IL-27 receptor, revealed increased activation of CD4^+^ T cells and IFN-γ production [32]. This enhanced immune response improved macrophage function and reduced bacterial burden. However, this beneficial effect was counterbalanced by a more severe chronic inflammatory state, ultimately accelerating death in WSX-1 KO mice. Supporting the proposed mechanism, the significant decrease in IL-27 levels in male B cell IL-10 KO mice observed in our study coincided with increased IFN-γ production, reduced bacterial burden, and enhanced inflammation, mainly due to a rise in CD68^+^ macrophages. Importantly, despite the increased inflammation, survival rates in these male B cell IL-10 KO mice were extended compared to their wild-type littermates. Presumably, the inflammation was still regulated to some extent and did not progress to the severe, damaging characteristics typically observed in complete absence of IL-27. These findings strongly suggest that B cell-derived IL-10 plays a critical, previously unrecognized role in determining the sex-specific immune response during TB infection. Moreover, this is the first study to demonstrate the potential regulation of IL-27 by IL-10. Whether the reduction in IL-27 expression in males is a direct consequence of B cell IL-10 deficiency or occurs indirectly via modulation of IFN-γ or other pathways warrants further investigation.

Sex disparities in immune responses are well-established, with females generally displaying a stronger humoral and cellular response and a more pro-inflammatory gene expression profile [33]. Previous studies have demonstrated that estrogen can regulate the IFN-γ promoter, leading to increased IFN-γ production [34, 35]. This suggests a potential role for estrogen in driving higher IFN-γ production in females during *Mtb* infection. However, a key question remains: why does B cell IL-10 specifically regulate IFN-γ expression in males but not females? We did not observe significant differences in the frequency of B10 cells between male and female mice. Hence, sex-specific regulation likely occurs downstream of IL-10 production. Interestingly, Islam et al. [36] observed increased STAT3 activation in male leukocytes in response to IL-10, which could potentially explain the specific suppression of IFN-γ in males through a pathway involving STAT3, known to downregulate IFN-γ [37]. Noteworthy, the JAK-STAT signaling pathway was one of the pathways most influenced by B cell IL-10 deficiency in males. However, the precise details of how B cell-derived IL-10 specifically activates STAT3 signaling only in males and how this impacts disease outcome requires further investigation.

While the JAK-STAT signaling pathway was downregulated in B cell IL-10 KO males, the host defense peptide pathway was enhanced. This, along with our histological data showing increased CD68^+^ macrophages in the lungs of these B cell IL-10 KO males, suggests a possible shift in the immune response towards a more macrophage-mediated defense mechanism in the absence of B cell-derived IL-10 in males. Recent research has revealed that tissue-resident B cells play a crucial role in regulating macrophage polarization and function, partly through the production of IL-10 [38]. This aligns with findings from a model of LPS-induced acute lung injury, where B cell-derived IL-10 inhibited macrophage activation and recruitment [39]. However, this has not been shown in the context of *Mtb* infection before, where studies have rather focused on the negative impact of T cell or macrophage-derived IL-10 on macrophage polarization and function [40–42].

Our study uncovers a previously unknown sex-specific role for B cell-derived IL-10 in TB immunity. The observation of enhanced STAT3 activation in males compared to females [36] hints at a potential factor contributing to this sex-specific effect. Moreover, possible reasons for the observed sex disparity in B cell-derived IL-10 regulation might include differences in the location, timing, and overall vigor of the resulting immune responses. Sex hormones could potentially influence the localization of B cells producing IL-10, leading to distinct patterns of immune regulation in males and females. This concept aligns with existing research demonstrating the suppression of B cell positioning and germinal-center formation by testosterone [43]. Alterations in B cell distribution in the *Mtb* infected lungs, as evidenced by our previous observation of smaller B cell follicles in males compared to females [6, 7], suggests B cells might interact very differently with other immune cells within the infected tissue, potentially influencing the production and localization of IL-10 and thus shaping the overall immune response in a sex-specific manner. Additionally, the timing of IL-10 production by B cells might be modulated by sex, impacting the initial stages of the immune response differently in males and females. Furthermore, variations in IL-10 receptor expression could be another contributing factor. Differences in receptor availability might lead to distinct responses to IL-10 signaling between males and females, potentially explaining the variations observed in the suppressive effect on immune responses.

Finally, another critical question remains: would these effects be mirrored by T cell-derived IL-10, or does B cell IL-10 have distinct, non-redundant functions during TB in males? A study by Madan et al. [22] demonstrating the non-redundant role of B cell-derived IL-10 in suppressing virus-specific CD8 T cell responses during cytomegalovirus infection highlights the potential for such unique functions. While the negative impact of T cell-derived IL-10 on TB immunity is well-established, these studies were primarily conducted in female animals. This raises the intriguing possibility that B cell IL-10 serves a distinct, sex-specific regulatory role due to potential differences in localization or timing between male and female B cell responses.

In conclusion, our findings highlight a role for B cell IL-10 in regulating sex differences in the immune response observed in TB infection. Understanding these sex-based disparities in B cell distribution and IL-10 regulation could be crucial for developing more targeted and effective TB treatment strategies.

## Acknowledgements

We would like to thank Silvia Maass for culturing *Mtb*; Kathleen Kurwahn (ISEF Lübeck) and the staff of the animal facility at the Research Center Borstel for animal care; Christian Rosero and Jasmin Tiebach for their help with the immunohistochemistry and preparation of the NanoString cartridge and Martina Hein for her help with the flow cytometry experiments. In addition, we would like to express our gratitude to Can Pinar and Marie Krüger of Cytolytics for their assistance in analyzing our flow cytometry data using the Cytolution software. This work was supported by the Life Science-Stiftung (FZB 2020.01) and in-house funding from the Research Center Borstel.

## Author contributions

B.E.S. and D.H. conceived and designed the experiments; D.H., S.M., L.E. and L.v.B. performed the experimental work; D.H., S.M., G.H.P. and J.B. analyzed the data; D.D.J. processed and scanned the Nanostring cartridge; J.B., T.G., and R.M. provided advice, mice and reagents; B.E.S. and D.H. wrote the paper. All authors revised the manuscript.

## Additional information

Correspondence to B.E.S.

## References

1. WHO, Global tuberculosis report 2023. Geneva: World Health Organization; 2023. Licence: CC BY-NC-SA 3.0 IGO. 2023.

2. Borgdorff, M.W., et al., Gender and tuberculosis: a comparison of prevalence surveys with notification data to explore sex differences in case detection. Int J Tuberc Lung Dis, 2000. 4(2):p. 123–32.

3. Hertz, D. and B. Schneider, Sex differences in tuberculosis. Semin Immunopathol, 2019. 41(2): p. 225–237.

4. Horton, K.C., et al., Sex Differences in Tuberculosis Burden and Notifications in Low- and Middle-Income Countries: A Systematic Review and Meta-analysis. PLoS Med, 2016. 13(9): p. e1002119.

5. Gupta, M., et al., Genetic and hormonal mechanisms underlying sex-specific immune responses in tuberculosis. Trends Immunol, 2022. 43(8): p. 640–656.

6. Dibbern, J., L. Eggers, and B.E. Schneider, Sex differences in the C57BL/6 model of Mycobacterium tuberculosis infection. Sci Rep, 2017. 7(1): p. 10957.

7. Hertz, D., et al., Increased male susceptibility to Mycobacterium tuberculosis infection is associated with smaller B cell follicles in the lungs. Sci Rep, 2020. 10(1): p. 5142.

8. Khader, S.A., et al., IL-23 is required for long-term control of Mycobacterium tuberculosis and B cell follicle formation in the infected lung. J Immunol, 2011. 187(10): p. 5402–7.

9. Slight, S.R., et al., CXCR5(+) T helper cells mediate protective immunity against tuberculosis. J Clin Invest, 2013. 123(2): p. 712–26.

10. Ulrichs, T., et al., Differential organization of the local immune response in patients with active cavitary tuberculosis or with nonprogressive tuberculoma. J Infect Dis, 2005. 192(1): p. 89–97.

11. Ulrichs, T., et al., Human tuberculous granulomas induce peripheral lymphoid follicle-like structures to orchestrate local host defence in the lung. J Pathol, 2004. 204(2): p. 217–28.

12. Linge, I., E. Kondratieva, and A. Apt, Prolonged B-Lymphocyte-Mediated Immune and Inflammatory Responses to Tuberculosis Infection in the Lungs of TB-Resistant Mice. Int J Mol Sci, 2023. 24(2).

13. Swanson, R.V., et al., Antigen-specific B cells direct T follicular-like helper cells into lymphoid follicles to mediate Mycobacterium tuberculosis control. Nat Immunol, 2023. 24(5): p. 855–868.

14. Choreno-Parra, J.A., et al., Mycobacterium tuberculosis HN878 Infection Induces Human-Like B-Cell Follicles in Mice. J Infect Dis, 2020. 221(10): p. 1636–1646.

15. Maglione, P.J., J. Xu, and J. Chan, B cells moderate inflammatory progression and enhance bacterial containment upon pulmonary challenge with Mycobacterium tuberculosis. J Immunol, 2007. 178(11): p. 7222–34.

16. Tedder, T.F., B10 cells: a functionally defined regulatory B cell subset. J Immunol, 2015. 194(4): p. 1395–401.

17. Redford, P.S., P.J. Murray, and A. O’Garra, The role of IL-10 in immune regulation during M. tuberculosis infection. Mucosal Immunol, 2011. 4(3): p. 261–70.

18. Moreira-Teixeira, L., et al., T Cell-Derived IL-10 Impairs Host Resistance to Mycobacterium tuberculosis Infection. J Immunol, 2017. 199(2): p. 613–623.

19. Yuan, C., et al., Mycobacterium tuberculosis Mannose-Capped Lipoarabinomannan Induces IL-10-Producing B Cells and Hinders CD4(+)Th1 Immunity. iScience, 2019. 11: p. 13–30.

20. du Plessis, W.J., et al., The Functional Response of B Cells to Antigenic Stimulation: A Preliminary Report of Latent Tuberculosis. PLoS One, 2016. 11(4): p. e0152710.

21. du Plessis, W.J., G. Walzl, and A.G. Loxton, B cells as multi-functional players during Mycobacterium tuberculosis infection and disease. Tuberculosis (Edinb), 2016. 97: p. 118–25.

22. Madan, R., et al., Nonredundant roles for B cell-derived IL-10 in immune counter-regulation. J Immunol, 2009. 183(4): p. 2312–20.

23. Meng, L., et al., Bone Marrow Plasma Cells Modulate Local Myeloid-Lineage Differentiation via IL-10. Front Immunol, 2019. 10: p. 1183.

24. Bankhead, P., et al., QuPath: Open source software for digital pathology image analysis. Sci Rep, 2017. 7(1): p. 16878.

25. Johnson, K., Create Reports Using Phenoptics Data. phenoptrReports, 2022 Available online: https://akoyabio.github.io/phenoptrReports/.

26. Luckey, D., K. Medina, and V. Taneja, B cells as effectors and regulators of sex-biased arthritis. Autoimmunity, 2012. 45(5): p. 364–76.

27. Fillatreau, S., Natural regulatory plasma cells. Curr Opin Immunol, 2018. 55: p. 62–66.

28. Ritter, K., J. Rousseau, and C. Holscher, Interleukin-27 in Tuberculosis: A Sheep in Wolf’s Clothing? Front Immunol, 2021. 12: p. 810602.

29. Cooper, A.M., et al., IFN-gamma and NO in mycobacterial disease: new jobs for old hands. Trends Microbiol, 2002. 10(5): p. 221–6.

30. Cooper, A.M., et al., Disseminated tuberculosis in interferon gamma gene-disrupted mice. J Exp Med, 1993. 178(6): p. 2243–7.

31. Erdmann, H., et al., The increased protection and pathology in Mycobacterium tuberculosis-infected IL-27R-alpha-deficient mice is supported by IL-17A and is associated with the IL-17A-induced expansion of multifunctional T cells. Mucosal Immunol, 2018. 11(4): p. 1168–1180.

32. Holscher, C., et al., The IL-27 receptor chain WSX-1 differentially regulates antibacterial immunity and survival during experimental tuberculosis. J Immunol, 2005. 174(6): p. 3534–44.

33. Klein, S.L. and K.L. Flanagan, Sex differences in immune responses. Nat Rev Immunol, 2016. 16(10): p. 626–38.

34. Fox, H.S., B.L. Bond, and T.G. Parslow, Estrogen regulates the IFN-gamma promoter. J Immunol, 1991. 146(12): p. 4362–7.

35. Brown, M.A. and M.A. Su, An Inconvenient Variable: Sex Hormones and Their Impact on T Cell Responses. J Immunol, 2019. 202(7): p. 1927–1933.

36. Islam, H., et al., Sex differences in IL-10’s anti-inflammatory function: greater STAT3 activation and stronger inhibition of TNF-alpha production in male blood leukocytes ex vivo. Am J Physiol Cell Physiol, 2022. 322(6): p. C1095–C1104.

37. Wan, C.K., et al., Opposing roles of STAT1 and STAT3 in IL-21 function in CD4+ T cells. Proc Natl Acad Sci U S A, 2015. 112(30): p. 9394–9.

38. Suchanek, O., et al., Tissue-resident B cells orchestrate macrophage polarisation and function. Nat Commun, 2023. 14(1): p. 7081.

39. Sun, Z., et al., B cell-derived IL-10 promotes the resolution of lipopolysaccharide-induced acute lung injury. Cell Death Dis, 2023. 14(7): p. 418.

40. Murray, P.J., et al., T cell-derived IL-10 antagonizes macrophage function in mycobacterial infection. J Immunol, 1997. 158(1): p. 315–21.

41. Schreiber, T., et al., Autocrine IL-10 induces hallmarks of alternative activation in macrophages and suppresses antituberculosis effector mechanisms without compromising T cell immunity. J Immunol, 2009. 183(2): p. 1301–12.

42. Ring, S., et al., Blocking IL-10 receptor signaling ameliorates Mycobacterium tuberculosis infection during influenza-induced exacerbation. JCI Insight, 2019. 5(10).

43. Zhao, R., et al., A GPR174-CCL21 module imparts sexual dimorphism to humoral immunity. Nature, 2020. 577(7790): p. 416–420.

