## Supplemental Figure S1 for "Ablation of B cell-derived IL-10 increases tuberculosis resistance"

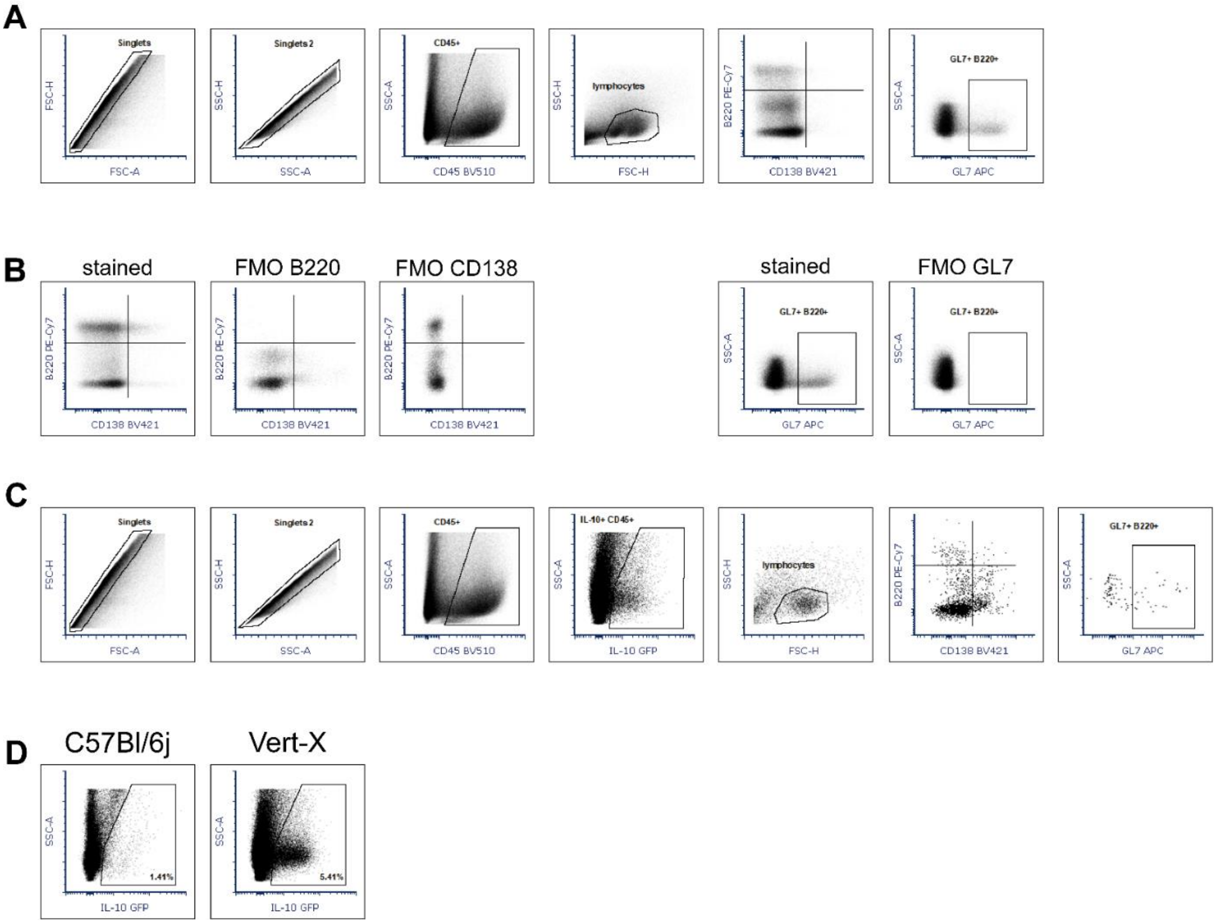


**Figure S1**. *Gating strategy to analyze IL-10 expression in B cells*. Representative flow cytometry plots for detection of B cell subtypes (A) using FMOs for B220, CD138, and GL7 (B) and the expression of IL-10 by B cell subtypes (C). An uninfected wild type C57BL/6j mouse was used as negative control for IL-10 expression (D).
