## Supplemental Figure S2 for "Ablation of B cell-derived IL-10 increases tuberculosis resistance"

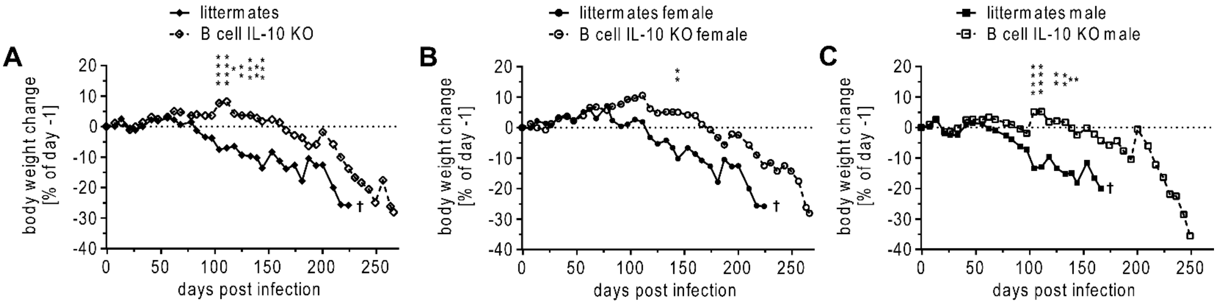


**Figure S2**. *Body weight changes of Mtb HN878 infected B cell IL-10 KO mice and littermates*. A) Body weight change (%) of B cell IL-10 KO mice (n = 18) and B cell IL-10 competent littermates (n = 12). Data stratified by sex for (B) females (n = 9 B cell IL-10 KO; n = 5 littermates) and (C) males (n = 9 B cell IL-10 KO; n = 7 littermates). Data represent the mean body weight change of animals from one experiment. Statistical analysis was performed by 2way ANOVA followed by Tukey's multiple comparisons test. *p<0.05; **p<0.01; ***p<0.001; ****p<0.0001
