## Supplemental Table S1 for "Ablation of B cell-derived IL-10 increases tuberculosis resistance"

| **Cycle** | **1** | **2** | **3** | **4** | **4** | **Nuclei stain** |
|---|---|---|---|---|---|---|
| **Antibody target** | Neutrophile Elastase | CD68 | CD3 | CD19 | | DAPI |
| **Clonality or clone** | polyclonal | polyclonal | SP7 | EPR23 174-145 | | |
| **Vendor, article number** | Abcam, ab68672 | Abcam, ab125212 | Zytomed Systems,RBK024 | Abcam, ab245235 | | |
| **Host** | rabbit | rabbit | rabbit | rabbit | | |
| **AR [pH6/pH9];** time [min] @ microwave power [Watt] | pH6;1min @1000, 10 min@ 100 | pH6;1min @1000, 10 min@ 100 | pH6;1min @1000, 10 min@ 100 | pH6;1min @1000, 10 min@ 100 | pH6; 1min @1000, 5 min@ 100 | - |
| 3% $H_2O_2$ Block | 0 | 0 | 0 | 0 | 0 | 0 |
| 1x TBST washing [min.] | 2, 2, 2 | 2, 2, 2 | pause over night | 2, 2, 2 | 2, 2, 2 | 2, 2, 2 |
| Protein blocking | 10 min. | 10 min. | 10 min. | 10 min. | OPAL 780 | Spectral DAPI |
| 1x TBST washing [min.] | 2, 2, 2 | 2, 2, 2 | 2, 2, 2 | 2, 2, 2 | | |
| Dilution | 1/100 | 1/250 | 1/200 | 1/200 | 1/25 | 3 drops in 1000µl PBS |
| time [min] | **45 min.** | **45 min.** | **45 min.** | **45 min.** | **60 min.** | **5 min**. |
| temperature[RT/4°C] | RT | RT | RT | RT | RT | RT |
| 1x TBST washing [min.] | 3, 3, 3 | 3, 3, 3 | 3, 3, 3 | 3, 3, 3 | 3, 3, 3 | 3, 3, 3 |
| Opal HRP Polymer dilution & [min.]@RT | 1/2, 10 min. | 1/4, 10 min. | 1/3, 10 min. | 1/4, 10 min. | Wash buffer | $dH_2O$ rinsing |
| 1x TBST washing [min.] | 3, 3, 3 | 3, 3, 3 | 3, 3, 3 | 3, 3, 3 | 3, 3, 3 | |
| **OPAL fluorophore & dilution** | **480, 1/150** | **520, 1/150** | **570, 1/150** | **OPAL TSA-DIG, 1/100** | Wash buffer | Mount with Prolong Gold |
| time [min] @ RT | 10 min. | 10 min. | 10 min. | 10 min. | 10 min. | |

**Table S1.** *Pipetting protocol for mIF staining.* Antibody targets and staining protocol to investigate immune cell populations within the infected lung of female and male B cell IL-10 KO mice and their littermates using mIF.
