## Supplemental Table S2 for "Ablation of B cell-derived IL-10 increases tuberculosis resistance"

| |
|---|
| body weight day before infection [g] |
| body weight experimental day [g] |
| CFU/lung |
| CFU/spleen |
| affected lung area [%] |
| B cell area [% of total lung area) |
| $CD19^+$ B cells [count/$mm^2$ lung area] |
| $CD3^+$ T cells [count/$mm^2$ lung area] |
| $CD68^+$ macrophages [count/$mm^2$ lung area] |
| $NE^+$ neutrophils [count/$mm^2$ lung area] |
| cytokine levels [pg/ml] |
| chemokine levels [pg/ml] |

**Table S2.** *List for PCA.* The various data obtained for female (n=6) and male (n=3) B cell IL-10 KO mice and their female (n=3) and male (n=5) B cell IL-10 competent littermates at day 82 after infection were used to generate a PCA plot.
